# Allometric correlates of heat tolerance in birds: A test using quail breeds with extreme size variation

**DOI:** 10.64898/2026.05.15.725344

**Authors:** Elin Persson, Joshua K. R. Tabh, Josefin Svensson, Andreas Nord

## Abstract

Birds and mammals are shrinking and shapeshifting as global temperatures rise. Ecogeographic rules predict that such changes should ease heat stress by increasing surface-area-to-volume ratios, and thus, the capacity for heat exchange. This has led to the hypothesis that body size reductions are driven by thermoregulatory selection or adaptive plasticity, although recent syntheses point to more complex, multifactorial causes. Crucially, recent theoretical models predict that thermoregulatory benefits of smaller body size only emerge at extreme deviations from average phenotypes. Here, we exploit agricultural selection in Japanese quail to directly test this hypothesis, using three breeds spanning extreme differences in body mass, surface area, and relative appendage lengths. Evaporative cooling capacity and the scope for evaporative water loss broadly followed allometric predictions when contrasting small and larger breeds. As expected, this allowed the smallest breed to tolerate higher air temperatures. However, differences in heat tolerance limits between breeds were consistently much smaller than predicted. Additionally, the breadth of thermoneutral zones overlapped in full, and upper critical temperatures were remarkably similar, between breeds. Together, these results show that heat tolerance is only weakly linked to surface-area-to-volume relationships and cannot be explained by size alone. Thus, although smaller bodies may modestly enhance heat dissipation when size variation in a population is substantial, our findings suggest that recent body size reductions and morphological shifts are unlikely to be driven primarily by thermoregulatory benefits.

## Introduction

Evidence shows that numerous mammals and birds have become smaller and shifted shape over the last several decades (Ryding et al., 2021; Searing et al., 2023; Sheridan & Bickford, 2011), with mean decadal declines of around 1% (Docampo et al., 2019; Weeks et al., 2020; Youngflesh et al., 2022). These changes coincide with climate warming-related increases in global temperatures and largely mirror expectations from Bergmann’s (Bergmann, 1847) and Allen’s (Allen, 1877) rules – that body size decreases and appendage length increases nearer the warm equatorial regions than in the cool polar regions. Hence, it has been proposed that size shrinkage reflects adaptation to alleviate thermoregulatory pressures in a warming world by increasing surface-area-to-volume ratios and therefore easing heat dissipation.

While the mechanisms driving this shrinkage may be complex (Baldwin et al., 2022; Martins et al., 2023; Shipley et al., 2022), a rich body of literature suggests that plastic responses to variations in developmental temperature may well be at play in both birds and mammals (Andrew et al., 2017; Burness et al., 2013; Cunningham et al., 2013; Demicka & Caputa, 1993; Rodriguez & Barba, 2016; van de Ven et al., 2020; see Weeks et al., 2022; Tabh & Nord, 2023 for reviews). Beyond this, selection for adult morphologies with high heat dissipation propensity may also contribute to size reductions and shape changes in a warming world, at least according to classic theory (*sensu* Mayr, 1956). While this second, selective mechanism has received considerable attention (e.g. Van Buskirk et al., 2010; Prokosch et al., 2019), it has also been historically questioned, given the extensive possibilities for endotherms to physiologically compensate for effects of size and shape on heat loss (e.g., by adjusting rates of evaporation, heat production, and superficial circulation and vasomotor state, using counter-current heat exchangers; Scholander et al. 1955; see also Steudel et al. 1994). This doubt is further supported by biophysical modelling suggesting that extant size declines in birds contribute, at most, minor direct thermoregulatory and thermolytic benefits (Nord et al., 2024). In line with this, we recently combined empirical measurements and predictive modelling to show that thermoregulatory benefits of a having a small body size in the heat were small and highly context-dependant, requiring larger deviations from standard, morphological allometries than what is typically present in natural populations (Tabh et al. 2025).

In this study, we exploited the outcomes of agricultural selection to empirically test the hypothesis that body size only influences thermoregulatory competence in the heat when individual phenotypes fall outside of the typical range for a species (Fig. 1A). We reared Japanese quail (*Coturnix japonica*) with three distinct body sizes (henceforth, ‘breeds; Fig. 1B) from hatching to adulthood, and measured heat tolerance limits (i.e., the highest air temperature a bird can tolerate without loss of motor control; Nord et al. 2026) and thermoregulatory competence at this limit (metabolic rate, evaporative cooling capacity, evaporative scope) both at peak growth rate and at maturity. If heat tolerance is determined by body size alone through its effect on surface-area-to-volume ratios (i.e., physiological compensation for varying surface areas is limited or negligible), we would expect evaporative cooling capacity, a predictor of the heat tolerance limit (Persson et al. 2026), and evaporative scope to scale curvilinearly with differences in surface-area-to-volume ratios between the breeds (Fig. 1C). By contrast, if body size itself, even when varying substantially, has no influence on heat tolerance limits and thermoregulatory competence (i.e., complete, physiological compensation is possible), we expected evaporative cooling capacity and scope to scale independent of body size. Finally, if heat tolerance is determined by a combination of size and physiology, we expect partial compensation with scaling deviating from that between surface-area-to-volume and breed.

**Figure 1.**
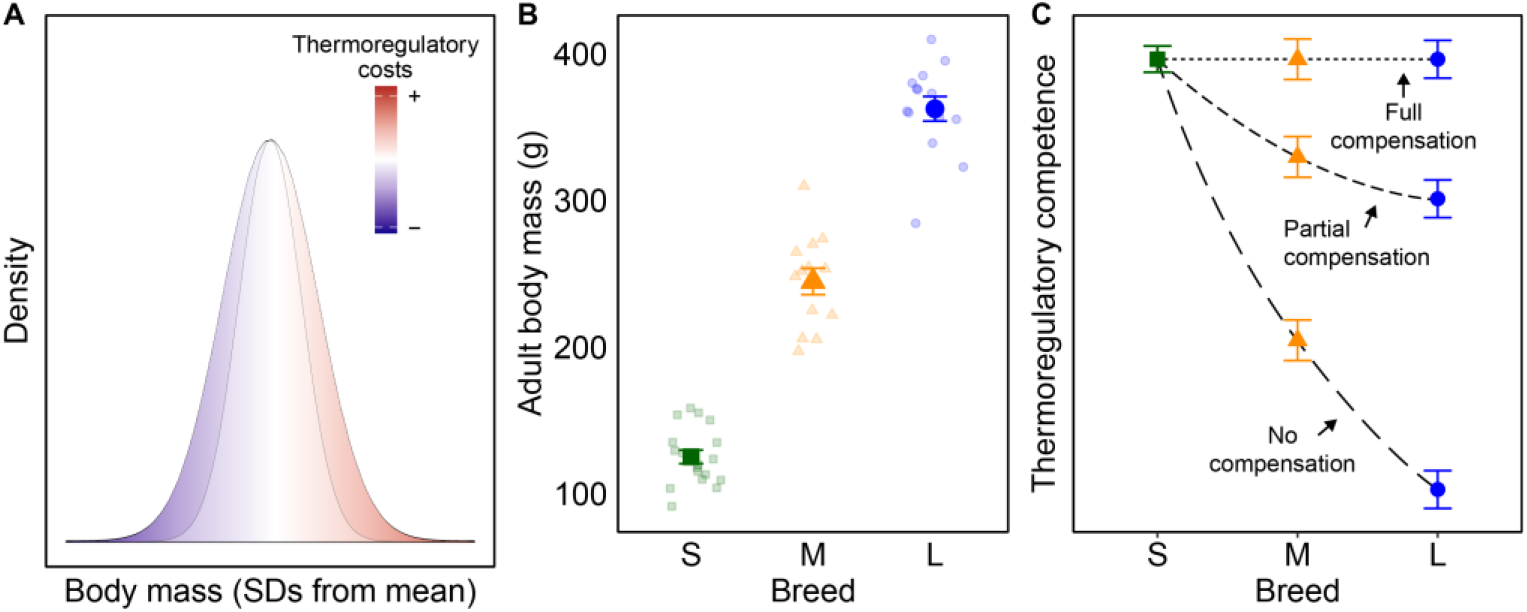
Conceptual model describing the effects of body size on thermoregulatory competence during heat exposure. A) Previous research indicates that costs of heat exposure only manifest when body dimensions deviate from standard allometries to an extent that is rarely observed in nature (Tabh et al., 2025). B) We harnessed outcomes of selection for agriculture to test this effect, using three breeds of Japanese quail with varying body masses. C) If body size alone determines heat tolerance, we expect thermoregulatory competence traits (e.g., evaporative cooling capacity, evaporative scope) to scale with differences in surface-area-to-volume ratios between the breeds (‘No compensation’), here predicted from body mass using published allometries (Walsberg & King, 1978). If, on the other hand, thermoregulatory physiology can compensate fully for the biophysical effect of size on heat flux (cf. Scholander, 1955), we expect a slope of zero (‘Full compensation’). In a situation where it is the combination of morphological and physiological traits that ultimately determine heat tolerance, we expect the performance-to-breed slope to diverge from pure size-effect slopes (‘Partial compensation’).

## Methods

### Species and husbandry

Japanese quail eggs from one small breed and one medium-sized breed were bought from commercial breeders. Eggs from one large breed were collected from an in-house population. While it is possible that these breeds may have experienced varying degrees of domestication, how this may influence different thermo-physiological traits is unclear. Nevertheless, by using three breeds along a size gradient, effects arising from differences in surface-area-to-volume ratio may be isolated from any domestication effects, if present. All eggs were kept at room temperature (18-20°C) in an automatic egg turner that turned the eggs for 5 min every 1 h until the start of incubation. In total, one hundred and thirty-seven eggs (n_small_ = 55; n_medium_ = 35; n_large_ = 46) were incubated in a Brinsea Ova Easy 190 incubator (Brinsea, Weston-super-Mare, United Kingdom) at 37.5°C and 50% relative humidity (regulated by a Brinsea Ova-Easy Advance Humidity Pump). Egg mass differed significantly between size categories (small: 9.4 ± 0.6 g; medium: 12.2 ± 0.7 g; large: 14.0 ± 0.5 g [mean ± s.e.]; *p* < 0.0001; Table S1).

At day 16 of incubation, eggs were moved from the incubation trays to hatching trays with individual compartments for each egg. We checked the eggs for hatching twice daily from day 17. Seventy-one eggs (n_small_ = 27; n_medium_ = 16; n_large_ = 28) hatched after an incubation period of 18 days (range: 17-19 days; mean ± s.d.: 17.92 ± 0.47 days). There was no difference in hatching time between the three size categories (Anova: *F* = 1.36, d.f. = 2,68, *p* = 0.2632), and average hatching success (≈52%, marginalised across size categories) was comparable to that observed in previous studies (Ben-Ezra & Burness, 2017; Persson et al., 2024). Seven birds died of natural causes, or were euthanized due to irreversible disabilities, during the first days after hatch. Of the remaining birds, 48 (n_small_ = 18; n_medium_ = 13; n_large_ = 17) were used in the experiments.

Chicks were banded with uniquely numbered leg bands 1 to 14 h after hatching (i.e., when completely dry) and were housed in breed-specific groups in open pens (155×120×60 cm; length × width × height) or in stacked cages (185×105×55 cm) lined with wood shavings. Birds had ad libitum access to food, water, seashells and sand. Until 3 weeks of age, the chicks were fed turkey starter (Kalkonfoder Start, Lantmännen, Stockholm, Sweden; 25.5% protein). From 3 weeks onwards, birds were fed turkey grower (Kalkonfoder Tillväxt, Lantmännen, Stockholm, Sweden; 22.5% protein). Mealworms, shredded carrots, or a combination of both, were provided as daily diet enrichment. Vitamin supplements (Protect Vitamin, Lantmännen, Stockholm, Sweden) were added to the enrichment feed. During the first 2 weeks after hatch, to compensate for the absence of a brooding female, birds were provided access to a heat lamp (37-39°C at ground level).

### Recording morphology across breeds

Birds were weighed once weekly (± 0.1 g) from hatching to adulthood. Hatching body mass differed between groups (small: 6.1 ± 0.37 g; medium: 8.3 ± 0.41 g; large: 10.5 ± 0.27 g; Table S1, Fig. S1A). At about halfway to sexual maturity (3 weeks; Fig. S3), body mass had increased to 63.0 ± 4.4 g, 120.9 ± 4.8 g and 196.4 ± 3.2 g, in small, medium, and large, birds, respectively (Table S1; Fig. S1B). Wing lengths (measured ± 0.5 mm) at 3 weeks followed a similar differential pattern, being, on average, 88.3 ± 11.2 mm, 94.5 ± 1.3 mm and 105.0 ± 0.9, respectively. However, inflection points for body mass growth (i.e., the point in time where growth was the fastest) were similar between size categories, being 2.3 ± 0.02 (mean ± s.e.) weeks for small birds, 2.4 ± 0.02 weeks for medium birds and 2.3 ± 0.01 weeks for large birds (Table S3). Small birds subsequently reached an asymptotic mass of 126.4 ± 11.1 g, medium birds of 247.7 ± 8.5 g, and large birds of 383.7 ± 11.4 g (Fig. S3), and had at 10 weeks of age wing lengths of 110.6 ± 1.0 mm, 120.3 ± 1.1 mm and 126.1 ± 0.7 mm in small, medium, and large, birds (Table S1; Fig. S1C). The birds reached 90% of asymptotic mass at 7.0, 6.3 and 6.2 weeks, respectively. Growth curves and analyses thereof are detailed in the ESM (Table S3, Fig. S3). Additionally, we used a combination of digital photography and modelling to analyse differences in surface-area-to-volume ratios, and relative appendage lengths, across breeds and ages. Detailed methodology and results are reported in the ESM (Table S2, Fig. S2).

### Measurement of heat tolerance limits and associated thermoregulatory traits

To measure and contrast heat tolerance characteristics between size categories, we measured metabolic rate, water loss, and body temperature responses to a heat ramp in a subsample of 43 birds (n_small_ = 15; n_medium_ = 13; n_large_ = 15). Subsampling was used to roughly balance representation among size categories. To allow for non-contact measurements of body temperature during metabolic measurements, we implanted a temperature-sensitive passive integrated transponder (< 0.5% of body weight; LifeChip BioTherm, Destron Fearing, South St. Paul, MN, USA) into the intraperitoneal cavity when the birds were 14 days old following Persson et al. (2024).

Heat tolerance measurements were performed using flow-through respirometry when the birds were 3 weeks old (range: 21-25 days; the period of most rapid growth) and 10 weeks old (range: 70-73 days; i.e., when asymptotic body size and reproductive maturity had been reached). Measurements were taken inside a climate-controlled test chamber (Weiss Umwelttechnik C180, Reiskirchen, Germany) and were conducted during daytime. Briefly, birds were placed in a 13 L glass chamber ventilated with dry (drierite) atmospheric air (small: 12.1 ± 0.04 L/min [mean ± s.d.], standard temperature and pressure, dry [STPD]; medium: 15.0 ± 0.04 L/min; large: 15.0 ± 0.03 L/min), measured with an Alicat 20SLPM mass flow meter (Alicat Scientific Inc., Tucson, AZ, USA). We subsampled 398.1 ± 0.49 ml/min (mean ± s.d.; SS4 pump, Sable Systems, Las Vegas, NV, USA) for gas analyses, from which we measured water vapour density (RH300 analyser; Sable Systems), carbon dioxide production (CA-10 analyser; Sable Systems) and oxygen consumption (FC-10 analyser; Sable Systems). Prior to measuring oxygen consumption, water vapour and carbon dioxide were removed from the airstream (using drierite and ascarite, respectively). To remove the influence of faeces on evaporation, birds rested on a metal grid platform over a 1 – 2 cm layer of mineral oil (Whitfield et al., 2015). Birds were fasted with water ad libitum at room temperature for 0.5 to 2.3 h before the start of respirometry. Then, they were acclimated in the respirometry chambers at 30°C for 30 min until gas traces were stable before the start of the experiment. After acclimation, we collected at least 5 min of stable gas data at 30°C, after which air temperature was acutely increased to 40°C, and then in 2°C increments once gas traces had been stable for at least 5 min (Noakes et al., 2016; Talbot et al., 2018). All experiments started and ended with a recording of reference air (baseline). Air temperature was manually adjusted and measured at floor and ceiling levels inside the chambers using thermocouples (copper-constantan 36-gauge type T; TC-2000 thermocouple box; Sable systems). An experiment ended when body temperature increased above 45°C, or when evaporative water loss did not increase further for an increase in air temperature.

To bring body temperature down quickly once experiments ended, the ventral plumage was wetted with EtOH (70%) and the birds were placed in front of a fan (in a 52 × 32 × 30 cm cage), with water ad libitum for 20 min.

### Inferring thermoneutral zones based on metabolic rate and evaporation

Beyond metabolic and evaporative responses at the heat tolerance limit, we also investigated how body size variations influenced thermoregulation across a broader range of ecologically relevant air temperatures (5°C-37.5°C), by estimating lower and upper critical temperatures (Scholander et al. 1950) from metabolic rate and evaporation data. Forty-eight birds (n_small_ = 18; n_medium_ = 13; n_large_ = 17) of which 43 had previously been used in the heat tolerance measurements, were used to study how metabolic rate, evaporative water loss, and body temperature, were affected by air temperature. The measurements were performed during nighttime when at 13 weeks of age (range: 85-98 days old). Four birds were measured sequentially using the same equipment as in the heat tolerance measurements described above, but drierite scrubbers were not used to dry the influent airstream (see below for data handling). Birds were placed in 6 (small) or 9 L (medium and large) glass respirometry chambers ventilated with pressurised air (small: 6.1 ± 0.15 L/min [mean ± s.d.], STPD; medium: 8.3 ± 1.08 L/min; large: 8.8 ± 0.62 L/min; subsampling: 389.7 ± 3.38 ml/min, STPD) at 18:30 ± 5 min, and were exposed to either a warming program (i.e., a heat ramp; n_small_ = 9; n_medium_ = 6; n_large_ = 9) or a cooling program (i.e., a temperature decline; n_small_ = 9; n_medium_ = 7; n_large_ = 8). Details on the temperature acclimation and temperature ramping are described in Table S5 and Fig. S6. Briefly both programs started with a 1 h acclimation at 20°C, after which temperature was changed ± 5°C twice during 2 successive hours. In the final step of acclimation, temperature was either acutely decreased to 2.5°C (from 10°C; warming program) or acutely increased to 37.5°C (from 30°C; cooling program) and data collection started. In the warming program, temperature was first increased to 5°C, then incrementally by 5°C until 35°C, and finally, increased to 37.5°C in a single step. In the cooling program, temperature was decreased following the same temperature steps as in the warming program. Hence, the warming program finished at 37.5°C and the cooling program at 2.5°C. Each temperature cycle lasted 60 min, starting and ending with a 10 min baseline, with 10 min of sequential data per bird between these periods. Birds were weighed and returned to their cages after the program was finished in the morning.

Ethical approval was granted by the Malmö/Lund Animal Ethics Committee (permit no. 19735-22).

### Respirometry data handling

All respirometry data handling were performed using macros written for ExpeData (v. 1.9.27; Sable Systems). We defined an individual’s heat tolerance limit as the air temperature where evaporative water loss had reached its maximum. We defined maximum evaporative water loss as the most stable 3 min at this temperature, and also extracted the corresponding metabolic heat production data. Resting metabolic rate data and resting evaporative water loss from the nighttime measurements were extracted from the period corresponding to the most stable 2 min of oxygen consumption in each air temperature.

All equations used for gas analyses were taken from Lighton (2019). We mathematically dry-corrected carbon dioxide data using the water vapour and atmospheric pressure following Dalton’s law of partial pressures according to equation 8.7, before calculating carbon dioxide production (eq. 11.6). For data collected overnight, we also ‘dried’ the incurrent flow rate in an analogous manner (following equation 8.6). Oxygen consumption was calculated using eq. 11.1, and were converted to metabolic heat production (W), assuming 20 J per 1 ml of O2 (Kleiber, 1961). Evaporative water loss (g^−1^) was calculated using eq. 11.9. Evaporative cooling capacity was defined as the ratio between evaporative heat loss (W; converted from evaporative water loss, assuming it requires 2406 J to evaporate 1 ml of water [Wallace & Hobbs, 2006]) and metabolic heat production (Lasiewski et al., 1966). Evaporative scope was calculated as the fold-change between resting, thermoneutral, (i.e., at 30°C) and maximum (i.e., at the heat tolerance limit) evaporative water loss.

### Statistical analyses

#### Comparisons of heat tolerance limits and associated thermoregulatory traits

Statistical analyses and model diagnostics (e.g., residual checks) were performed using R (ver. 4.3.0; R Core Team, 2023). Differences in thermoregulatory traits between breeds, measured during heat tolerance trials, were analysed using linear mixed effects models (lmer() function in lme4; Bates et al., 2015). We used metabolic heat production (W; log-transformed to meet parametric assumptions), evaporative water loss (g^−1^), evaporative cooling capacity at the heat tolerance limit (unitless), the heat tolerance limit itself (°C), and evaporative scope (unitless) as response variables. Breed, age and breed×age were used as categorical explanatory variables, to test if any differences between sizes changed with age. Group-mean centred body mass (by age and breed) was used as a continuous covariate. To account for repeated measurements, bird identity was added as a random intercept.

To test if the response in metabolic heat production, evaporative water loss (log-transformed for 3-week data to meet parametric assumptions), evaporative cooling capacity, and body temperature, to increasing air temperature differed between size categories, we modelled metabolic heat production, evaporative cooling capacity and body temperature, against air temperature (continuous variable), breed and air temperature×breed. In models involving evaporative water loss, we also included temperature^2^ and temperature^2^×size as covariates since raw data indicated non-linearity. Group-mean centred body mass was used as a covariate, and bird identity as random intercept.

The significance of parameters from the linear mixed models were assessed with likelihood ratio tests. Non-significant interactions were removed from the models, but all main effects were kept. Differences between factor levels or significant interactions were compared using post hoc tests (pairs () in the emmeans package; Lenth, 2023). Back-transformed estimates of metabolic heat production and evaporative water loss are presented in the text and figure.

#### Determining thermoneutral zones using metabolic rate and evaporation data

To identify and ultimately compare limits of thermoneutrality across size categories, we used manually-constructed breakpoint regressions with metabolic heat production and evaporative water loss as response variables, and air temperature as the fixed effect. Here, we defined the breakpoint for metabolic heat production across air temperature as the lower critical temperature and the breakpoint for evaporative water loss as the upper critical temperature (IUPS Thermal Commission, 2003). We also defined the breakpoints for body temperature and evaporative cooling capacity.

To construct our breakpoint regressions, we first fitted size-specific linear mixed models (using the lme() from the nlme package; Pinheiro et al., 2025) with metabolic heat production, evaporative water loss, evaporative cooling capacity, or body temperature as response variables, measurement temperature as a continuous covariate, group-mean centred body mass as a covariate, and bird identity as a random intercept. These models were then extended with a segmented regression using segmented() from (segmented package; Muggeo, 2008) to estimate breakpoints and their confidence intervals (CIs). An initial starting value for the breakpoint of 25°C was used in each size group, deemed appropriate from visual inspection of the raw data.

## Results

### Small quail were more heat tolerant than medium and large quail

The small breed of Japanese quail had significantly higher heat tolerance limit (44.50 ± 0.346°C) than both the medium (42.40 ± 0.354°C) and large breeds (41.70 ± 0.325°C), exceeding each by 2.1 and 2.8°C respectively (Table 1; Fig. 2A).

**Table 1.**
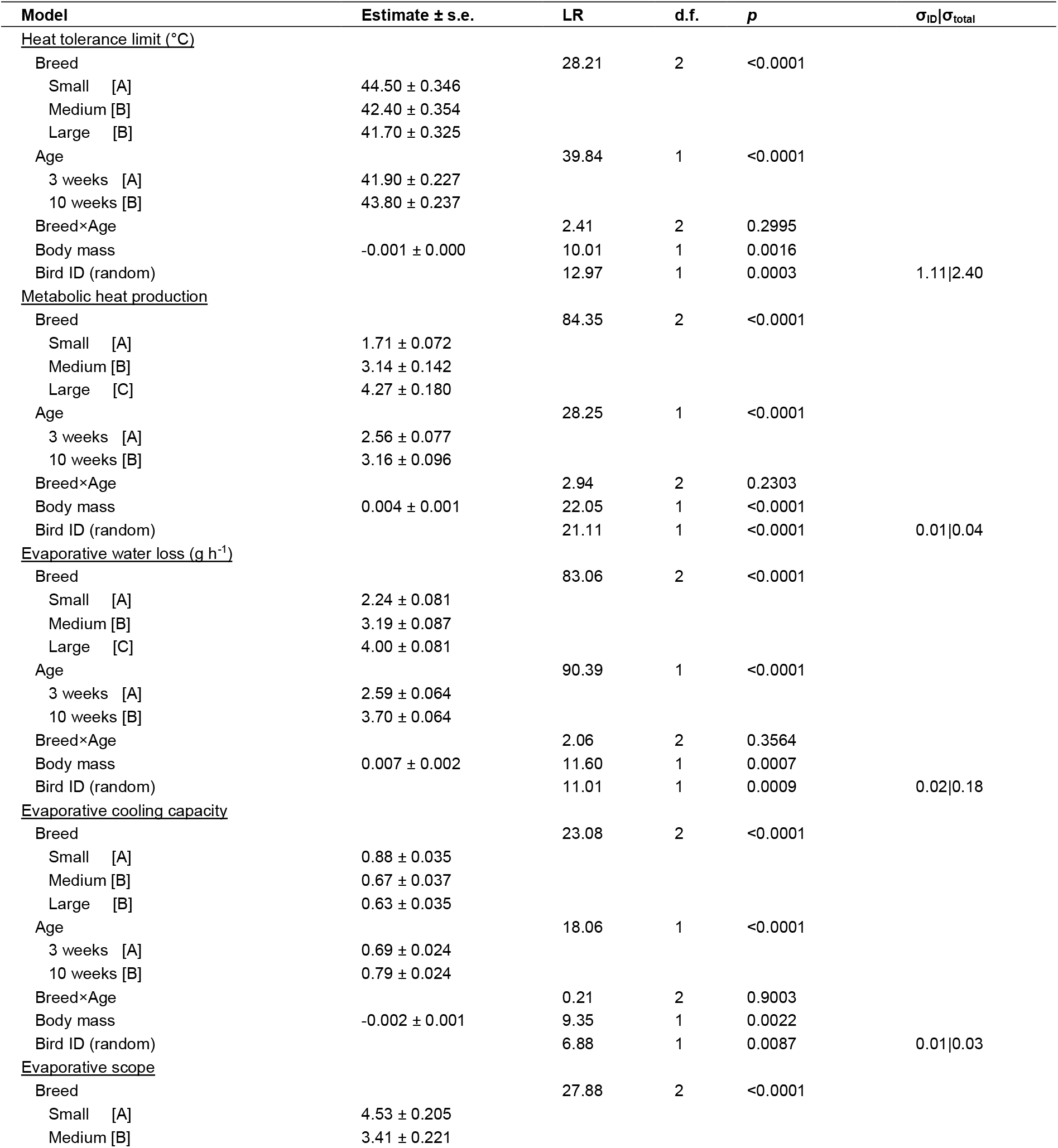
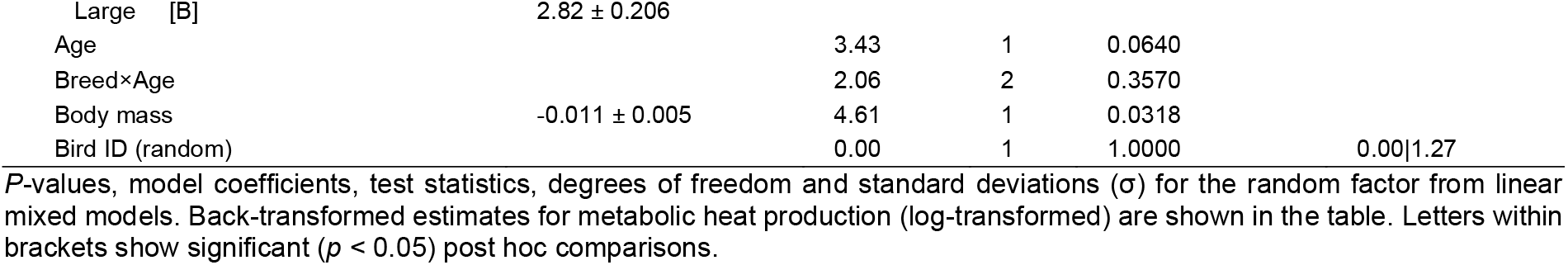
Parameter estimates from models evaluating the effects of body size across three breeds of Japanese quail on heat tolerance limit, metabolic heat production, evaporative water loss, evaporative cooling capacity, and evaporative scope at 3 and 10 weeks of age.

**Figure 2.**
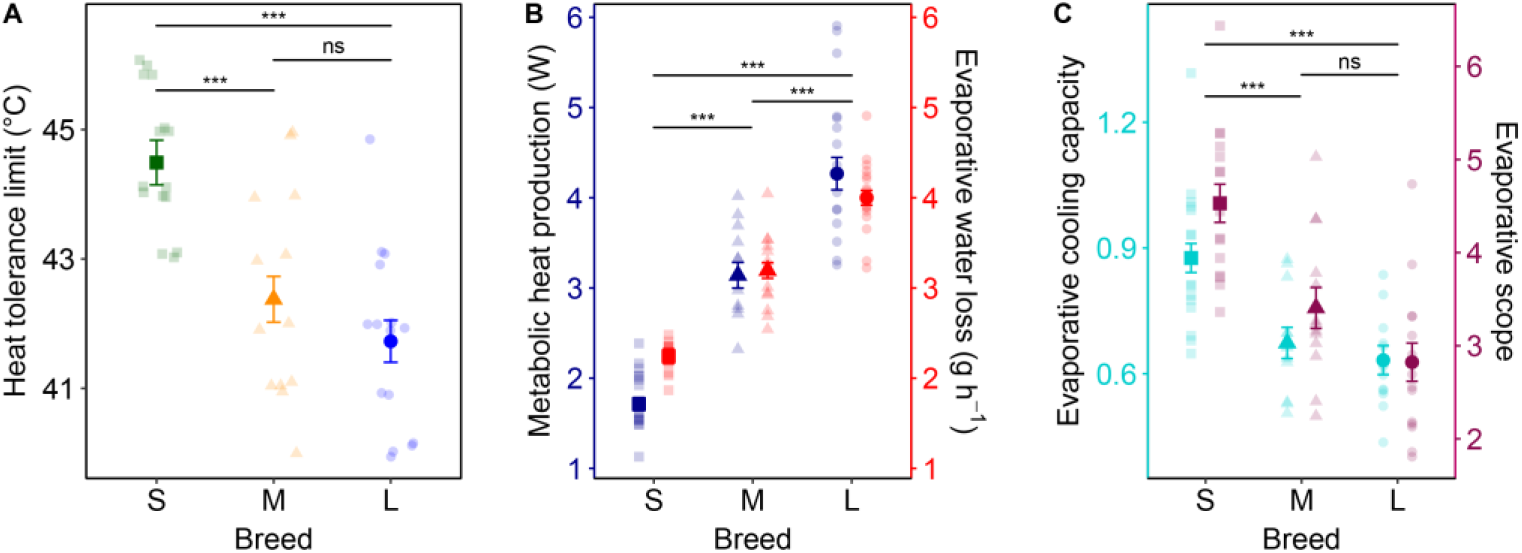
Thermoregulatory traits at the heat tolerance limit in small, medium and large quail breeds. Panels show (A) the heat tolerance limit – the highest air temperature at which the birds could maintain stable evaporation and avoid severe, damaging, hyperthermia. B) metabolic heat production and evaporative water loss at the heat tolerance limit; and C) evaporative cooling capacity (i.e., the ratio between metabolic heat production and evaporative heat loss) and evaporative scope (i.e., the fold-change from thermoneutral to maximal evaporation) at the tolerance limit. All data are averaged over 3- and 10-week-old birds. Points and error bars show estimated means ± 1.0 standard error (s.e.), and the semi-transparent points show the raw data means of age-specific measurements. Asterisks show significance levels (ns: *p* > 0.05; *: 0.05 ≥ *p* > 0.01; **: 0.01 ≥ *p* > 0.001; ***: *p* ≤ 0.001).

Both metabolic heat production and evaporative water loss scaled with body mass at the heat tolerance limit. Specifically, across ages, metabolic heat production was the lowest in the small breed (1.71 ± 0.072 W), followed by the medium and large breeds (3.14 ± 0.142 and 4.27 ± 0.180 W, respectively; Table 1; Fig. 2B). Evaporative water loss at the heat tolerance limit followed the same pattern, with the small breed evaporating the lowest amount of water per unit time (2.24 ± 0.081 g^−1^) followed by the medium (3.19 ± 0.087 g h^−1^) and large breeds (4.00 ± 0.081 g h^−1^; Table 1; Fig. 2B). However, the small breed displayed higher evaporative cooling capacity (0.88 ± 0.035) than both the medium (0.67 ± 0.037) and large breeds (0.63 ± 0.035; Table 1; Fig. 2C). Furthermore, the small breed had 1.3 to 1.6-fold higher evaporative scope (4.53 ± 0.205) than the others (medium: 3.41 ± 0.221; large: 2.82 ± 0.206; Table 1; Fig. 2C). The effect of size on all of these thermoregulatory traits was identical across both ages (all size × age: *p* > 0.05; Table 1). On average, however, 10-week-old birds had higher intercepts than 3-week-old birds in all traits except evaporative scope (that did not differ between ages; Table 1; Fig. S4).

Thermoregulatory responses to increasing air temperature – that is, the slope of trait-to-temperature relationships – differed between breeds (Table S4; Fig. S5). Specifically, the small breed did not increase metabolic heat production towards the heat tolerance limit at any age, whereas both the medium and large breeds did. Evaporative water loss, on the other hand, increased faster in the small and medium breeds compared with the large breed at 3 weeks, and with increasing size between breeds at 10 weeks. As a result, evaporative cooling capacity increased faster with air temperature in the small compared with the large breed at 3 weeks, and faster in the small breed compared with the others at 10 weeks (Fig. S5C). The rate of increase in body temperature towards the heat tolerance limit, equalling 0.13°C per 1 °C increase in air temperature on average, did not differ between breeds, but large breed had higher body temperature intercepts than the small breed at 3 weeks (by ~0.4°C) and higher intercepts than both the small and medium breeds at 10 weeks (by ~0.3 – 0.5°C) Fig. S5D).

### No differences in thermoneutral zones between breeds

Based on metabolic heat production and evaporative water loss, the thermoneutral zone in the small breed was estimated to be between 22.9°C (95% CI 19.0 – 26.9°C; Fig. 3A) and 29.2°C (95% CI 28.2 – 30.3°C; Fig. 3A), and the body temperature breakpoint to 28.2°C (95% CI 26.9 – 29.6°C; Fig. 3B). In the medium-sized breed, the estimated thermoneutral zone was very narrow, ranging from 28.3°C (95% CI 25.6 – 31.0°C; Fig. 3A) to 28.7°C (95% CI 27.5 – 29.9°C; Fig. 3A), and the upper body temperature breakpoint was estimated at 27.9°C (95% CI 26.1 – 29.8°C; Fig. 3B). Similarly to the small breed, the thermoneutral zone in the largest breed ranged between 23.4°C (95% CI 20.8 – 25.9°C; Fig. 3A) and 29.0°C (95% CI 28.1 – 29.9°C; Fig. 3A), but the body temperature breakpoint was lower in the large when compared with the small and medium breeds (22.9°C; 95% CI 21.4 – 24.4°C; Fig. 3B). The air temperature of evaporative cooling capacity inflection points was estimated to 29.2°C in the small breed (95% CI 28.2 – 30.3°C), 27.9°C in the medium breed (95% CI 26.6 – 29.1°C), and 28.9°C in the large breed (95% CI 28.1 – 29.7°C).

**Figure 3.**
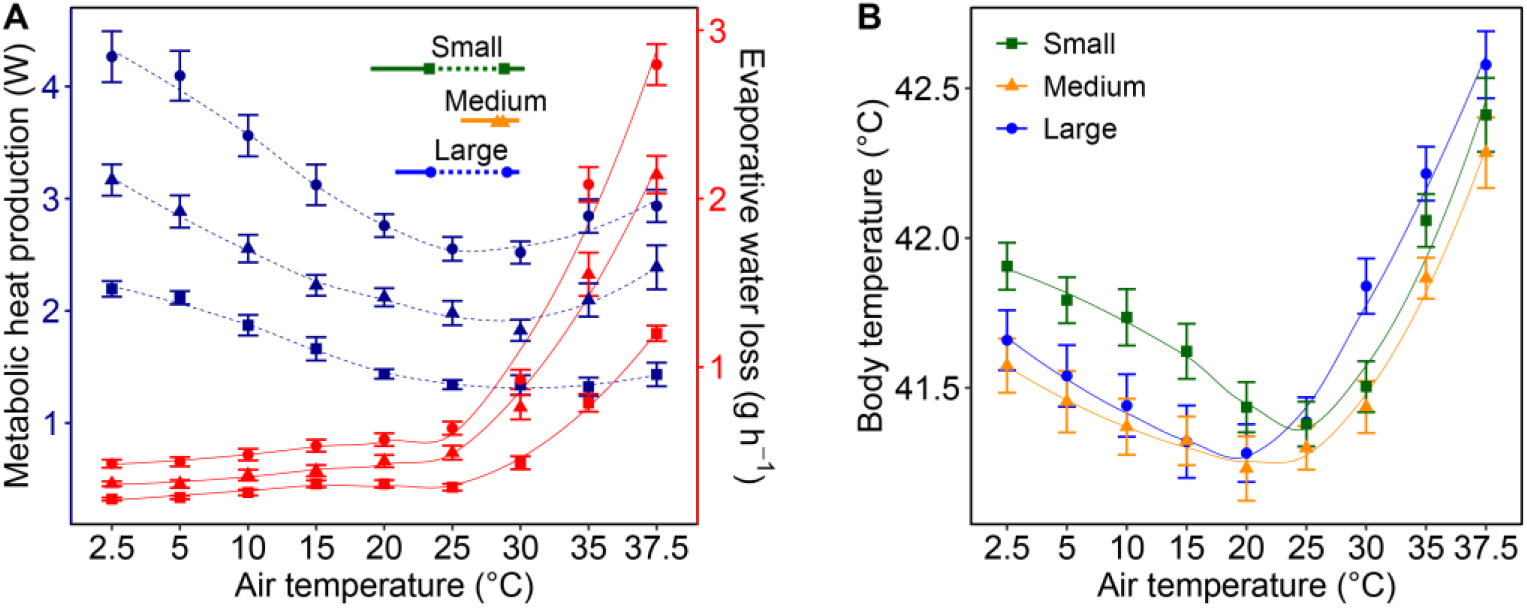
Thermoneutral zones and thermoregulatory breakpoints across size breeds of Japanese quail. A) Curves show metabolic heat production (blue) and evaporative water loss (red) across a range of ecologically relevant air temperatures. Curves with points and error bars show a LOESS smoother (locally estimated scatterplot smoothing) overlaid on raw data means ± 1.0 standard error (s.e.). The inset, horizontal, lines show breed-specific thermoneutral zones estimates, derived from breakpoint regressions of metabolic and evaporation data. The dotted lines show the estimated thermoneutral zone, solid points demarcate the lower and upper critical temperatures, and solid lines extend to 95% CIs (mean – CI, mean + CI). B) Means ± s.e. with LOESS smoothers showing body temperature as a function of air temperature for the differently sized quail breeds. All measurements depicted in the panels were collected during nighttime in resting and fasted animals.

## Discussion

In line with our predictions, the small breed of Japanese quail tolerated higher air temperatures than the medium and large breeds before reaching maximum evaporative capacity and facing body temperature dysregulation. This difference was correlated with shifts in evaporative cooling capacity and evaporative scope; traits reflecting thermoregulatory competence during heat exposure. Scaling followed – at least in part – expectations from variation in surface-area-to-volume ratios and relative appendage lengths. For instance, surface-area-to-volume ratios were 44% larger, and evaporative cooling capacity 39% higher, in the small compared with the large breed, supporting our ‘no compensation hypothesis’ (Fig. 1C). However, differences in evaporation traits between the medium and large breeds were below allometric expectations (‘partial compensation’). Additionally, the scaling of heat tolerance limits consistently fell below allometries (differing by 2 to 6% between breeds), thermoneutral zones overlapped almost fully and upper critical temperatures were remarkably similar. In all, these findings indicate that even when thermoregulatory traits follow allometric predictions, heat tolerance is only weakly linked to surface-area-to-volume relationships despite extreme size variation. This observation casts some doubts on the hypothesis that extant size reductions among avian lineages can be explained by selection or adaptive plasticity for thermoregulatory benefit alone.

Our results support recent predictive studies suggesting that morphologies must fall substantially outside population distributions to meaningfully alleviate thermoregulation during warm temperatures (Nord et al., 2024; Tabh et al., 2025). Even so, thermoregulatory benefits of a reduced body size in the heat are likely to vary by environmental, evolutionary, and physiological context.

First, the influence of surface-area-to-volume ratios on passive heat dissipation (i.e., via conduction, convection, and radiation) diminishes as environmental temperature approaches body temperature, and disappears at thermal equilibrium. Beyond this point, a smaller body size may even become disadvantageous, as it increases the surface area available for heat gain (e.g., Mitchell et al., 2018). Consistent with this, heat loss avenues at the heat tolerance limit (approximated using thermal conductance; Aschoff, 1981) in our study was almost entirely evaporative.

Second, skin surface area is comparatively less important for adaptively increasing evaporation during acute heat exposure. At mild or thermoneutral temperatures, evaporative water loss in birds is typically divided approximately equally between cutaneous and respiratory pathways, whereas at high environmental temperatures heat dissipation becomes dominated by respiratory mechanisms (Wolf & Walsberg, 1996; Wojciechowski et al., 2020). This reflects the fact that, in most bird species, cutaneous evaporation is not believed to be actively regulated over short timescales (but see Ro & Williams, 2011). Indeed, rapid, facultative increases in cutaneous evaporation have been convincingly demonstrated only in columbids (Marder & Gavrieli-Levin, 1986; 1987). Consequently, increases in relative skin surface area are unlikely to substantially enhance heat dissipation under hot conditions, compared with the more flexible and actively regulated respiratory pathways, with few exceptions. That said, allometry may still influence respiratory evaporation. For example, in quail, we recently found that individuals with larger bill surface areas increased evaporative water loss more rapidly during heat exposure (Tabh et al. 2025), potentially reflecting greater exposure of moist mucosal surfaces that enhanced the efficiency of gular fluttering.

Third, it may be an oversimplification to assume that heat tolerance limits themselves scale in proportion to changes in surface-area-to-volume ratios. Although thermolysis and thermogenesis are regulated via largely distinct pathways (Persson et al., 2026), they remain phenotypically correlated and cannot be fully uncoupled physiologically. This interdependence is likely to constrain the extent to which differences in body size can be compensated. Consistent with this, the medium- and large-bodied breeds in our study incurred measurable metabolic costs during heat exposure, whereas the small breed did not. At the same time, small birds exhibited lower body temperature intercepts and increased evaporative water loss more rapidly as temperatures approached the heat tolerance limit. Together, these patterns suggest that thermoregulatory control in small-bodied quail was both more effective and more flexible than in larger individuals. It remains to be determined to what extent this conclusion can be generalised across avian species and populations.

Our results do not refute the idea that smaller body size can enhance acute heat tolerance under some conditions, particularly when size differences exceed allometric expectations. However, larger body sizes may contrastingly confer advantages during prolonged heat exposure by reducing the risks of dehydration (McKechnie & Wolf, 2021) and starvation (Goodman et al., 2012). In addition, larger individuals may outcompete smaller ones for access to microhabitats, food, and water when resources are limited. These benefits, together with the well-established links between body size, fecundity, and reproductive fitness (e.g., Ronget et al., 2018) may offset – or even outweigh – the relatively modest thermoregulatory advantages of smaller size (cf. Siepielski et al., 2019). Consequently, understanding the role of body size declines in contemporary bird species requires a more integrative perspective that considers both the potential thermoregulatory benefits of small size and the ecological and physiological costs it may impose, particularly under conditions of resource limitation and water scarcity.

One implication arising from this work is that heat tolerance may be more usefully understood in terms of its consequences for the organism rather than as variation in thermoregulatory physiology alone. This is illustrated by the striking similarities in thermoneutral zone boundaries and upper critical temperatures among the breeds in our study, despite clear differences in heat tolerance. This mismatch aligns with expectations based on the shallow scaling of thermoneutral zone breadth in mammals (Riek & Geiser, 2013), and supports calls for caution when using metrics from the curves describing the interrelations between metabolic rate, evaporation, and air temperature (the ‘Scholander-Irving model’; Scholander, 1950) to predict heat tolerance in a warming world (McKechnie et al., 2016; Welman et al., 2024; Wolf et al., 2017). Together, these findings underscore the complexity of relationships between body size, physiology, and thermal limits. More specifically, they highlight the need to view heat tolerance holistically, as an integrated measure of the extent to which body size and shape enable birds to offset the cascading costs of heat exposure, from missed ecological opportunities (Cunningham et al., 2021) to cellular and organismal damage, including protein misfolding (Feder & Hofmann, 1999), oxidative stress (Belhadj Slimen et al., 2014), telomere attrition (Eastwood et al., 2022), mitochondrial dysfunction (Correia et al., 2025), impaired immunity (Oluwabenga & Fraley, 2023), and reduced gamete viability (Snook et al., 2026). Viewed in this light, future work may benefit from treating heat tolerance as an integrative outcome of these processes rather than as a single physiological threshold.

## Author contributions

Elin Persson, Joshua K. R. Tabh and Andreas Nord conceived the idea. Elin Persson and Andreas Nord designed the project. Elin Persson and Josefin Svensson performed the practical work, and data was analysed by Elin Persson, Andreas Nord and Joshua K. R. Tabh. Elin Persson and Andreas Nord wrote the first draft of the manuscript which was edited by all authors. Supervision was provided by Andreas Nord, and funding was procured by Andreas Nord, Elin Persson and Joshua K. R. Tabh.

## Acknowledgements

We thank Livia Saccani Hervas for help with bird care, and Lars Fredriksson and Kasper Hård for practical assistance.

## Ethics

Ethical approval was granted by the Malmö/Lund Animal Ethics Committee (permit no. 19735-22).

## Data availability

Data and code will be published upon acceptance in figshare.

